# Pancreatic α and β cells are globally phase-locked

**DOI:** 10.1101/2020.08.16.252304

**Authors:** Huixia Ren, Yanjun Li, Chengsheng Han, Yi Yu, Bowen Shi, Xiaohong Peng, Shufang Wu, Xiaojing Yang, Sneppen Kim, Liangyi Chen, Chao Tang

## Abstract

The Ca^2+^ modulated pulsatile secretion of glucagon and insulin by pancreatic α and β cells plays a key role in glucose homeostasis. However, how α and β cells coordinate via paracrine interaction to produce various Ca^2+^ oscillation patterns is still elusive. Using a microfluidic device and transgenic mice in which α and β cells were labeled with different colors, we were able to record islet Ca^2+^ signals at single cell level for long times. Upon glucose stimulation, we observed heterogeneous Ca^2+^ oscillation patterns intrinsic to each islet. After a transient period, the oscillations of α and β cells were globally phase-locked, i.e., the two types of cells in an islet each oscillate synchronously but with a phase shift between the two. While the activation of α cells displayed a fixed time delay of ~20 s to that of β cells, β cells activated with a tunable delay after the α cells. As a result, the tunable phase shift between α and β cells set the islet oscillation period and pattern. Furthermore, we demonstrated that the phase shift can be modulated by glucagon. A mathematical model of islet Ca^2+^ oscillation taking into consideration of the paracrine interaction was constructed, which quantitatively agreed with the experimental data. Our study highlights the importance of cell-cell interaction to generate stable but tunable islet oscillation patterns.

## INTRODUCTION

To precisely regulate the blood glucose level^1–3^, glucose elevation induces Ca^2+^ oscillation in pancreatic islet cells, which may trigger the pulsatile secretion of insulin and glucagon^4–7^. Dampening and disappearance of islet Ca^2+^ oscillation is an early biomarker in the pathogenesis of type 2 diabetes^8–11^. Multiple types of glucose-stimulated oscillation patterns have been observed in islets, including fast (~20 s cycle), slow (~100 s cycle), and mixed oscillations (20~100 s cycle)^12–14^. While different mathematical models have been proposed to explain the underlying mechanism^15–20^, most focused on the intrinsic properties of single or coupled β cells, such as the endoplasmic reticulum Ca^2+^ buffering capacity^18^ and the slow metabolic cycle of ATP/ADP ratio during glucose stimulation^17^. These β cell-centric models, however, may not fully explain the observed variety of oscillation patterns in islets. Only slow Ca^2+^ oscillations are mostly seen in isolated β cells^21–24^. Furthermore, glucagon accelerates the Ca^2+^ oscillation in isolated β cells^25^. The islet is a micro-organ in which multiple cell types closely interact. The α and β cells show highly correlated Ca^2+^ oscillation patterns^26^ and periodic release of insulin and glucagon is temporally coupled both *in vitro*^4,27^ and *in vivo*^6,7,28,29^. Thus the extensive autocrine and paracrine interactions between α and β cells^30–35^ may modulate or even dictate the islet oscillation modes.

The challenge to test such a hypothesis lies in resolving the identity of individual cells and monitoring their activity in live islets simultaneously. In addition, because the spatial organization of α and β cells are highly heterogeneous from islet to islet^36,37^, quantitative comparison of Ca^2+^ oscillations in different islets is necessary. To address these problems, we have developed a microfluidic device attached to the spinning-disc confocal microscope, which allowed individual cells to be imaged under physiological conditions for up to ~2 hrs. By the long-term imaging of islets undergoing repeated glucose stimulation, we found that the oscillation mode represents intrinsic properties of the islet. By constructing a new transgenic mice line, we could identify the cell types in live islets accurately. Quantitative analysis revealed generic features as well as quantitative relationships in the oscillation patterns across many islets. In particular, we found that oscillations of the islet α and β cells are each synchronized but phase shifted, and that the value of the phase shift between α and β cells determines the oscillation mode. Finally, we developed a coarse-grained mathematical model incorporating paracrine interactions between α and β cells. The model reproduced key quantitative features of the experimentally observed oscillations and suggested that different oscillation modes may come from the varied paracrine controls.

## RESULTS

### Glucose-Evoked Ca^2+^ Oscillation Represents an Intrinsic Property of the Islet

To provide a stable and controllable environment for long-term imaging of intact islets, we developed a microfluidic device (Fig. 1A). On one side, we designed an inlet port to load the islet (300 μm in width and 270 μm in height), which could be sealed after loading. The chip could trap islets of different sizes with a descending PDMS ceiling (270, 180, 150, 110, 80, and 50 μm in height). On the side opposite the inlet, five independent input channels merged into one channel upstream of the islet trapping site. Such a device enabled long-term and stable imaging of islets even during the switching of different perfusion solutions. For instance, when glucose concentration in the perfusion solution was increased from 3 to 10 mM (3G to 10G), all islets exhibited an initial rapid rise in cytosolic Ca^2+^, followed by a gradual appearance of fast (cycle<60 s, 31 of 46 islets), slow (cycle>60 s, 9 of 46 islets) or mixed (6 of 46 islets) Ca^2+^ oscillations (Fig. 1B). In contrast, although high glucose still evoked the initial increase in cytosolic Ca^2+^ concentration, no following periodic Ca^2+^ oscillations were observed in isolated islets from diabetic *db/db* mice (9 of 9 islets, Fig. 1B lower panel)^10,11^, suggesting the critical role of Ca^2+^ oscillation in maintaining normal hormone release and glucose homeostasis. While different islets displayed largely variable Ca^2+^ oscillations, the second round of 10G stimulation in the same islet evoked an oscillation frequency nearly identical to the first round (Fig. 1C). The spatial activation profiles were also similar, as almost identical cells lighted up at the designated times of Ca^2+^ cycles during the two rounds of stimulation (Fig. 1D). Quantitatively, sequential activations of islet cells between the two rounds of stimulation showed a significantly higher similarity index than random association (Fig. 1E, See Methods). Therefore, specific oscillation modes represent a robust intrinsic property of individual islets, possibly determined by the fixed spatial organization of the islet.

**Fig. 1.**
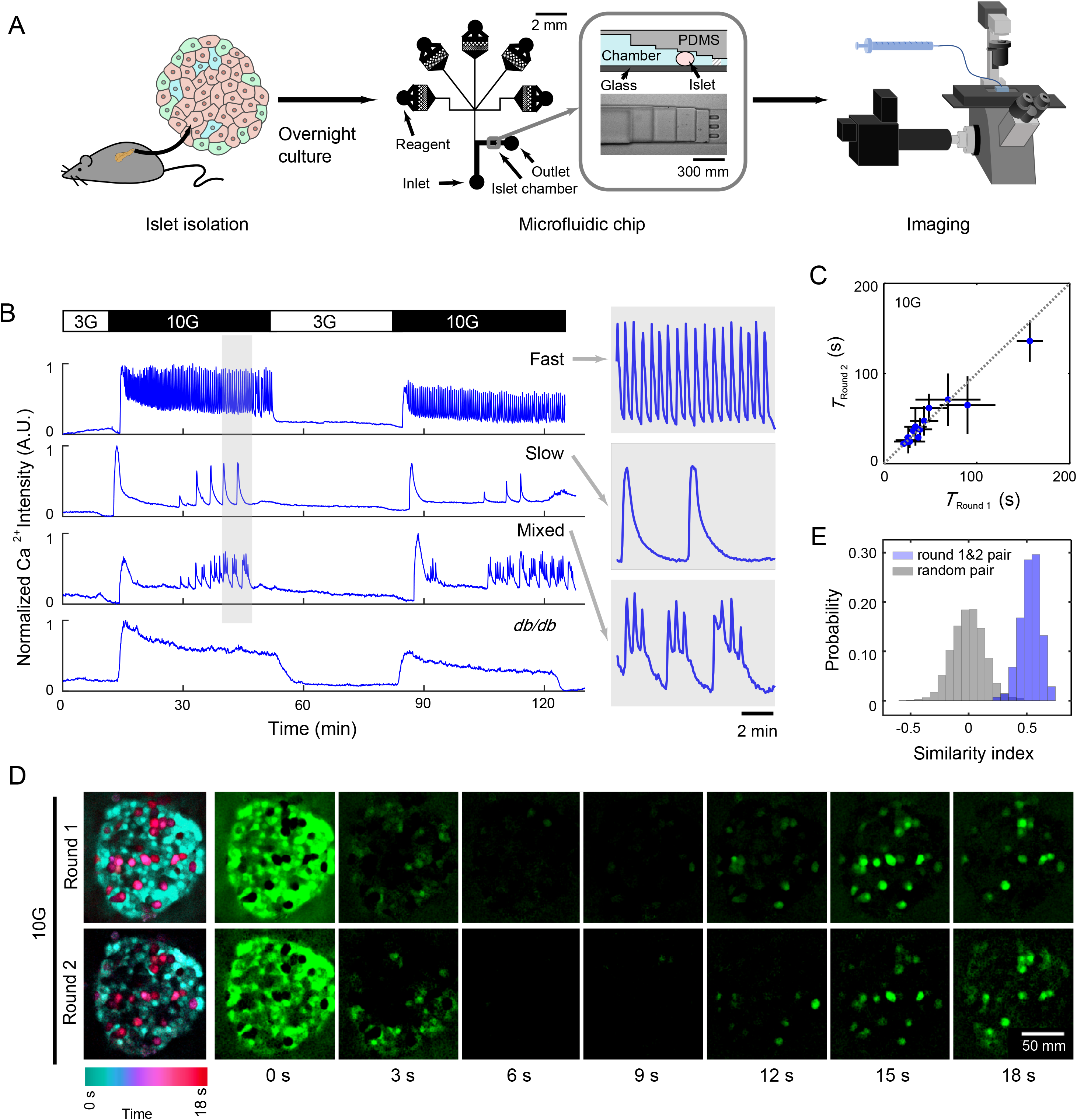
Islets Show Intrinsic Ca^2+^ Oscillation Modes under High Glucose Stimulation. (A) Experimental flow chart. Islets are isolated from mice. After overnight culture, the islets are loaded onto the microfluidic chip for imaging with confocal microscopy. The chip comprises five reagent channels, an inlet channel and an outlet channel. The islet chamber traps the islet with a gradient height from 270 μm to 50 μm. (B) Representative recordings of whole islet Ca^2+^ signal in Cal-520 AM loaded islets isolated from C57BL/6J mouse. The islet is stimulated with a repeated protocol: 10 min 3 mM glucose (3G), 40 min 10 mM glucose (10G), 30 min 3G and 40 min 10G. The first panel, islet displays fast oscillation with a period of ~20 s (31 of 46 islets); second panel, slow oscillation at ~3.5 min (9 of 46 islets); third panel, mixed oscillation at ~20 s and 2.45 min (6 of 46 islets); fourth panel, absence of Ca^2+^ oscillation in a *db/db* mouse islet (9 of 9 islets). Except for the last one, enlarged images of the shaded region are shown on the right. (C) The mean Ca^2+^ oscillation period during the first *versus* the second round of 10G stimulation (n = 24 islets). (D) Cell activation sequence in the first and second round of 10G stimulation. We subtract the previous frame from the next frame of the original Ca^2+^ images (frame interval 3 s). Shown is the cell activation sequence averaged across oscillation cycles in a 5 min interval (aligned with the maximum activation frame). Left panels summarize the time sequence shown in the 7 right panels, with the pseudo-color representing the activation time. (E) The same islet shows a high similarity index between the first and the second round of 10G stimulation (n=5 islets).

### Identification of Islet α and β Cell Types Using Transgenic Mice

To probe the specific micro-organization of an islet and to distinguish the Ca^2+^ activities between α and β cells, we generated *Glu-Cre^+^; GCaMP6f^f/+^; Ins2-RCaMP1.07* mice in which α and β cells were labeled with the green and red fluorescent Ca^2+^ sensor, GCaMP6f and RCaMP1.07, respectively (Fig. 2A, see Methods). Because the vector was randomly inserted into the genome using the PiggyBac transposon system, β cells were sparsely labeled (9.4%, Figs. 2B and S1). RCaMP1.07 is a Ca^2+^ sensor with a fluorescence on-rate similar to GCaMP6f^38^ and Cal-520 AM (Figs. S2C and S2D). We confirmed the labeling accuracy by immunofluorescence (Figs. 2C and S1). The RCaMP1.07 expressing islet cells were 100% insulin-positive, while the GCaMP6f expressing islet cells were 95.5% glucagon positive (Fig. S1A and S1B). This result was also confirmed in intact islets using immunofluorescence labeling (Figs. S1C and S1D) and pharmacology experiments (Figs. S2A and S2B). The GCaMP6f expressing cells responded to both NE and glutamate stimulation, while the RCaMP1.07 expressing cells were silent under both stimuli. These data both reinforced the expression specificity of α and β cells, and non-detectable overlaps in emission spectrums between GCaMP6f (525/50 nm) and RCaMP1.07 (600/50 nm) (Figs. 2D and S2E).

**Fig. 2.**
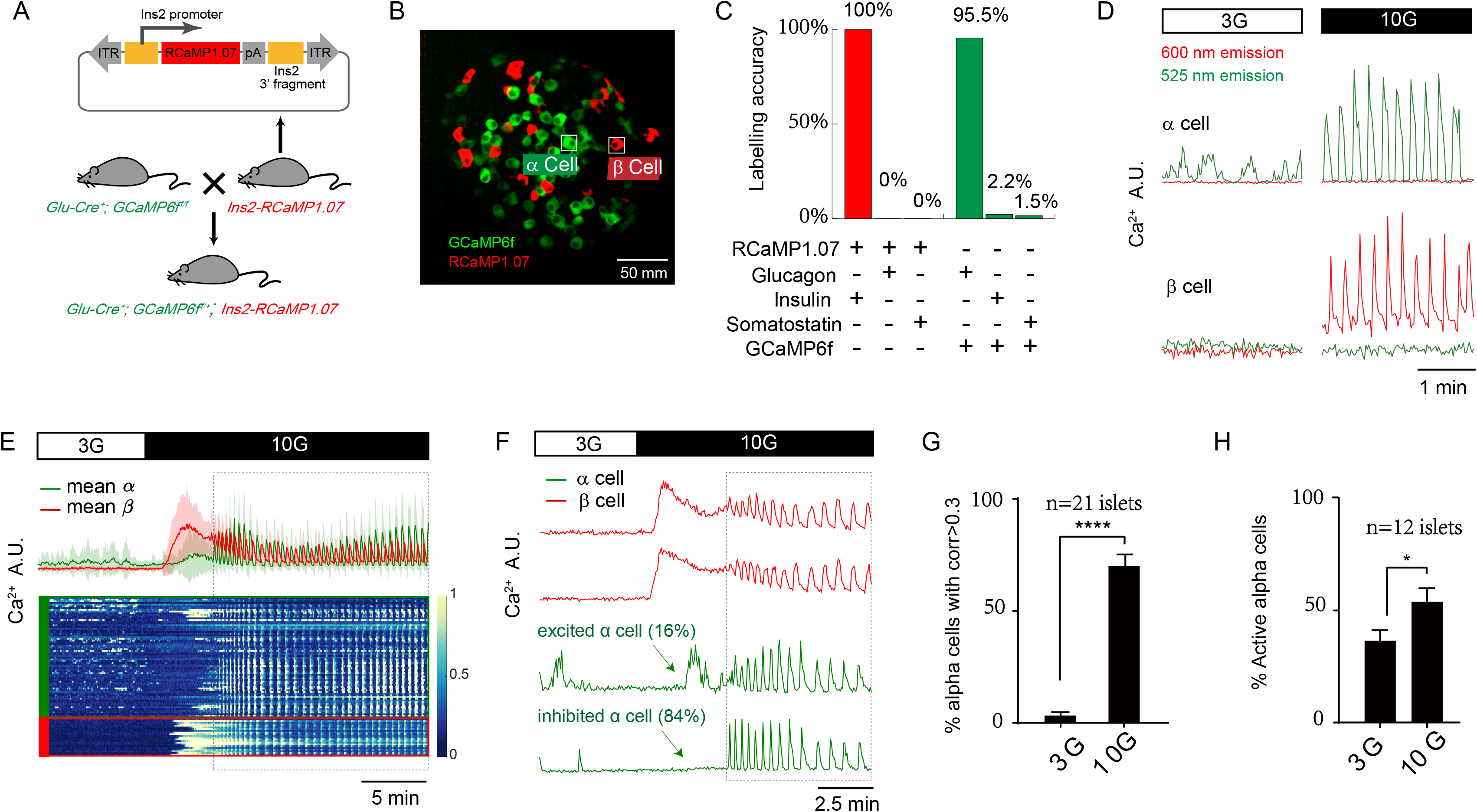
Using *Glu-Cre^+^; GCaMP6f^f/+^*; *Ins2*-*RCaMP1.07* transgenic Mice to Identify Islet α and β Cells. (A) Gene targeting vector designed with *Ins2* 5’-promoter, *RCaMP1.07* and *Ins2* 3’-fragment for the construction of the *Ins2-RCaMP1.07* mice. By crossbreeding *Ins2-RCaMP1.07* mice with *Glu-Cre^+^; GCaMP6f^f/f^* mice, we generate the *Glu-Cre^+^; GCaMP6f^f/+^*; *Ins2-RCaMP1.07* mice. (B) Maximal projection of Ca^2+^ activity from the *Glu-Cre^+^; GCaMP6f^f/+^*; *Ins2*-*RCaMP1.07* mice islet. α cells expressed GCaMP6f (Green) and β cells sparsely expressed RCaMP1.07 (Red) (see Methods). (C) Immunofluorescence co-localization analysis of *Ins2-RCaMP1.07* and *Glu-Cre^+^; GCaMP6f^f/f^* islet cells.100% RCaMP1.07 expressing cells are insulin positive (n=281 RCaMP1.07+ cells), 0% glucagon positive (n=10 RCaMP1.07+ cells) and 0% somatostatin positive (n=20 RCaMP1.07+ cells). GCaMP6f expressing cells are 95.5% glucagon positive (n=178 GCaMP6f+ cells), 2.2% insulin positive (n=90 GCaMP6f+ cells) and 1.5% (n=772 GCaMP6f+ cells) somatostatin positive. (D) 525 nm and 600 nm emission (single bandpass filter with width 50 nm) signals from single α and β cells under 3G (2.5 min) and 10G (2.5 min) stimulation. The cell positions are marked in (B). Note the GCaMP6f and RCaMP1.07 proteins are exclusively expressed in islet cells. (E) Mean α and β cells Ca^2+^ signal from intact mouse islets exposed to 3–10 mM glucose. Shading corresponds to s.d.. Lower panel is the heat-map of the normalized single-cell Ca^2+^ signal. Green labels are α-cells and red β-cells. Dashed Box shows the stable oscillatory phase. (F) Single α and β cell Ca^2+^ signal from intact mouse islets exposed to 3–10 mM glucose. Note that 16% α cells showed evoked Ca^2+^ response to glucose elevation and 84% were silent (n=179 α cells from 5 islets). Dashed Box shows the stable oscillatory phase. (G) Percentage of synchronized α cells (mean Pearson correlation coefficient >0.3) under 3G and 10G stimulations (n=21 islets). Note that α cells showed significantly higher correlation under high glucose stimulation. (H) Percentage of active α cells (normalized to 25 mM KCl stimulated α cell number) under 3G and 10G stimulations (n=12 islets).

Under resting glucose stimulation, β cells remained silent while α cells demonstrated variable Ca^2+^ transients (Fig. 2E, Video 1), agreeing with the previous reports^26^. Therefore, the mean Ca^2+^ transients of β cells were flat, while we noticed a fluctuated mean Ca^2+^ trace of α cells. Upon elevated glucose stimulation, β cells always responded with large initial rises in cytosolic Ca^2+^ followed by slow decays.

In contrast, α cells demonstrated two opposite Ca^2+^ responses (Fig. 2F): within the first five minutes after the 10G stimulation, the majority of α responder (84%) were significantly inhibited, while the remaining cells (16%) showed evoked Ca^2+^ transients (n=179 α cells from 5 islets). Combined with 25 mM KCl stimulation, we found that 10G activated more α cells than 3G (Fig. 2H). Although this minor population of α cells did not fit the consensus, glucose-stimulated Ca^2+^ responses in some α cells were also noted previously^39,40^. Therefore, there exist two types of α cells in intact islets.

### Islet α and β Cells Are Globally Phase-Locked

At the later stage of 10G stimulation, different pools of α cells in the islet became synchronized, accompanied by synchronized Ca^2+^ oscillations from β cells (Fig. 2E). Indeed, while less than 5% of α cells demonstrated correlated Ca^2+^ transients at the resting and the initial glucose stimulation stages, more than 70% became synchronized later (Figs. 2G, 3A and 3B, Video 2). Unlike β cells interconnected by the gap junction protein Connexin36 to achieve synchronization^41–45^, α cells do not express gap junction proteins and are thus not physically connected^46^. Because mean Ca^2+^ peaks of α cells displayed a fixed delay to those of β cells (~20 s), we hypothesized that the highly synchronous α cell activity might be due to stable phase-locking to the β cell activity (Figs. 3C and 3D). Consistent with tightly inhibited α cells originating from their neighboring β cells, we observed global α cell activation after the turning-off of β cells (Fig. 3E). Intriguingly, phase-locking with similar temporal characteristics was also present in slow and mixed oscillations (Fig. 3F, Videos 3 and 4), suggesting a common underlying mechanism.

**Fig. 3.**
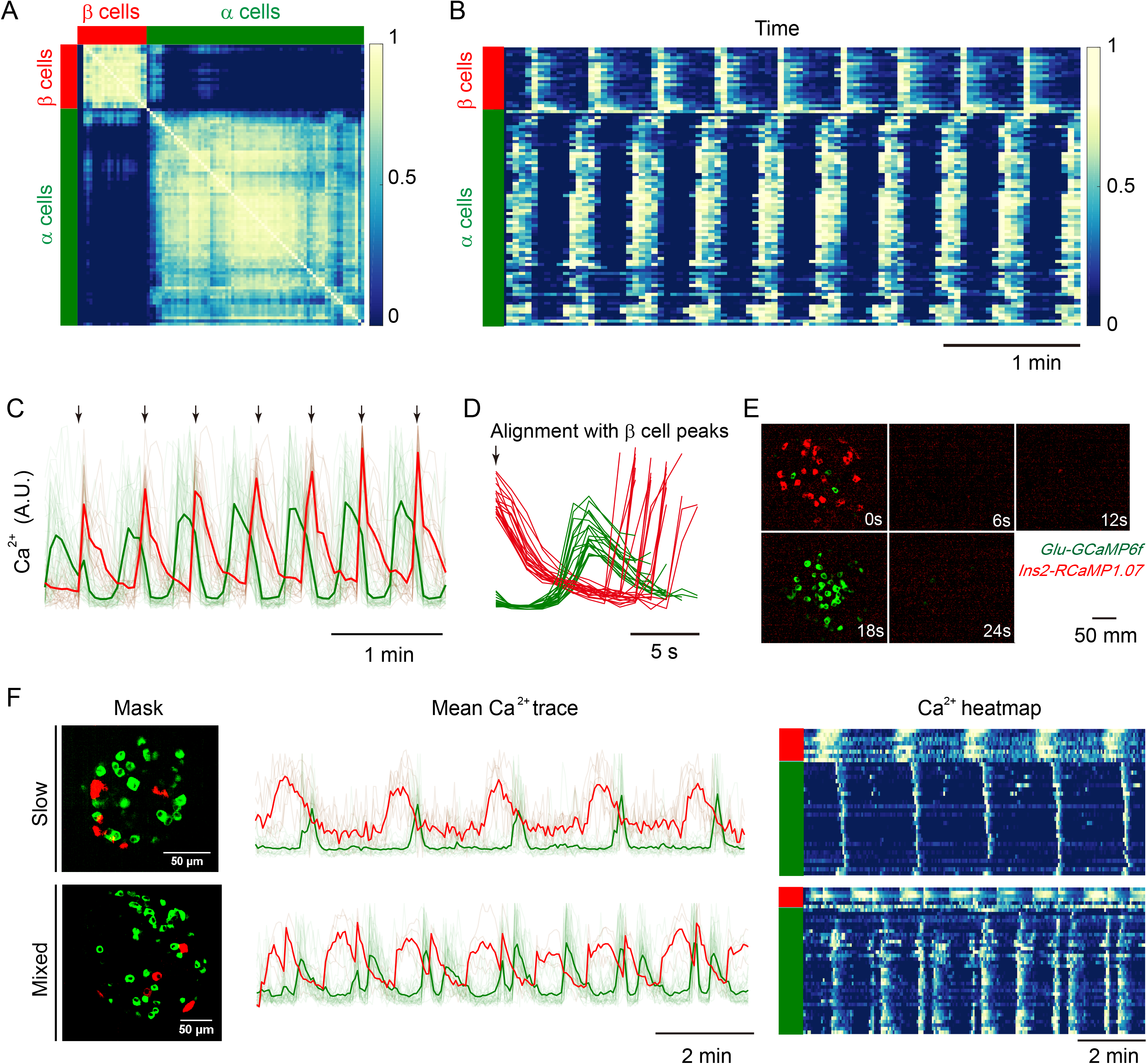
α and β Cells Are Globally Phase-Locked During Oscillation. (A) Correlation matrix (Pearson correlation coefficients) for Ca^2+^ activity of *Glu-Cre^+^;GCaMP6f^f/+^*; *Ins2* - *RCaMP1.07* islet cells under 10G stimulation. Cells are sorted according to cell types. β cells are indicated by red bar and α cells green bar. (B) The heat-map of time-dependent Ca^2+^ activity for α and β cells under 10G stimulation. Color bar codes the normalized Ca^2+^ intensity. β cells are indicated by red bar and α cells green bar. Each row represents the same cell in (A). (C) Mean Ca^2+^ activity of α and β cells under 10G stimulation for the same islet as A. Single cell traces are shown with light lines. The red trace represents β cells and green trace α cells (n=22 for β cells and n=71 for α cells). (D) Mean Ca^2+^ activity of α and β cells in C, aligned at each β cell peak. Each trace starts from a peak of β cell oscillation and stops in the next peak. (E) Sequential activation of α and β cells under 10G stimulation. We subtract the previous frame from the next frame of the original Ca^2+^ images and average across oscillations (each oscillation is aligned with the maximum activation frame). The mean activation sequence uses 5 min Ca^2+^ images (frame interval is 3 s). The β cells are colored red and α cells green. (F) Representative recordings of slow and mixed oscillations of Ca^2+^ activity of α and β cells under 10G stimulation. Top: slow oscillation; bottom: mixed oscillation. The first column, maximal projection of Ca^2+^ masks (α cells green and β cells red); second column, mean Ca^2+^ trace of α and β cells (single cell traces are shown with light lines); third column, heat-map of α and β cells’ Ca^2+^ activity. Color bar codes the normalized Ca^2+^ intensity.

The existence of phase-locked cells was also found in islets labeled with a non-targeted genetic Ca^2+^ sensor or loaded with the trappable Ca^2+^ dye (Video 5 and 6). In *Ella-Cre^+^; GCaMP6f^f/+^* islets in which all islet cells were labeled with *GCaMP6f*, islet cells could also be classified into two groups that are phase-locked (Video 5). In addition, in the *Glu-Cre^+^; tdTomato^f/+^* islet loaded with Cal-520 AM, we found that one of the two phase-locked cell populations colocalized with the genetically labeled red α cells (Video 6). Therefore, these data demonstrated that α cells are stably and globally phase-locked to β cells under elevated glucose stimulation in general.

### Variable Delay of β Cell Activation After α Cell Determines the Oscillation Mode

To non-biasedly sort out critical parameters differentiating various Ca^2+^ oscillation patterns, we quantitatively defined features of each oscillation cycle of alternately activated α and β cells (Figs. 4A and S3). The oscillation period *T* was defined as the time window between one cycle of β cell activation, which was split into two parts divided by the α cell activation: the waiting time for α cell to activate after β cell activation (*T*_βα_) and the waiting time for β cell to activate after α cell activation (*T*_αβ_). The phase difference (Δ*θ*) of two types of cells was defined as the ratio between *T*_βα_ and *T*. There were multiple ways to determine these quantities – either using the peak, 25%, 50%, or 75% decrease - and they did not change the outcome of the analysis (Figs. S4B-D). In total, we defined 13 features that recapitulated different characteristics of the α and β cell Ca^2+^ dynamics, such as the rise and decay time of the activation for α and β cells, the Full Width at Half Maximum (FWHM) (Table 1, and see Methods for detailed definitions). Based on these parameters, the oscillating cycles were vectorized and assembled into a feature matrix. Next, by dimension reduction of these parameters with the UMAP algorithm^47^, oscillations could be classified into the fast and the slow ones (Fig. 4B), in which *T*, Δ*θ*, *T*_αβ_ and FWHM of β cells demonstrated significant inter-group differences (Figs. 4C and S4A). The *T* for the fast oscillations centered at ~30 s, threefold smaller than the slow ones (~104 s). Similarly, the average Δ*θ* for the rapid oscillations was threefold larger than the slow ones (~0.6 versus ~0.2, Table 1). While the mean *T*_βα_ was almost indistinguishable between fast and slow oscillations, the *T*_αβ_ of the former was smaller than the latter (~11 s versus ~86 s, Table 1). The distribution of β cells’ FWHM for the fast oscillations was different from that of the slow ones, with a mean FWHM at ~12 s for the fast oscillations and ~34 s for the slow oscillations (Fig. 4C and Table 1). In contrast, the distributions of FWHM for α cells between the fast and slow oscillations were more similar (Fig. 4C), This implies a higher FWHM ratio of the Ca^2+^ transients of α to β cells in the fast Ca^2+^ oscillations (~1.02) than in the slow oscillations (~0.44) (Fig. S4A).

**Table 1.** The mean value of the features in the fast and slow clusters. Revised p value: p^r^ = p(×1E5). *p^r^ <0.05, **p^r^ <0.01, ***p^r^ <0.001, ****p^r^ <0.0001. Statistical comparisons were conducted using unpaired t test. Sample sizes of fast cluster and slow cluster were 574 pairs and 84 pairs, respectively. For detailed definitions, please see Methods.

| | Fast cluster | Slow cluster | p-value ($\times 1E5$) | Significance |
| --- | --- | --- | --- | --- |
| $T$ (s) | 29.8 | 99.9 | 3E-20 | **** |
| $\Delta\theta$ | 0.67 | 0.24 | 5E-111 | **** |
| $T_{\beta\alpha}$ (s) | 19.9 | 17.7 | 1.7 | n.s. |
| $T_{\alpha\beta}$ (s) | 9.5 | 81.2 | 2E-21 | **** |
| $\alpha_{FWHM}$ (s) | 11.9 | 14.8 | 0.08 | n.s. |
| $\beta_{FWHM}$ (s) | 11.4 | 31.1 | 8E-8 | **** |
| $\frac{\alpha_{FWHM}}{\beta_{FWHM}}$ | 1.04 | 0.48 | 4E-4 | *** |
| $\tau_{\alpha\_up}$ (s) | 5.1 | 5.0 | 7E-5 | n.s. |
| $\tau_{\alpha\_decay}$ (s) | 8.8 | 13.6 | 8E-7 | **** |
| $\tau_{\beta\_up}$ (s) | 5.1 | 18.7 | 7E-5 | **** |
| $\tau_{\beta\_decay}$ (s) | 10.4 | 20.4 | 2E-5 | **** |

**Fig. 4.**
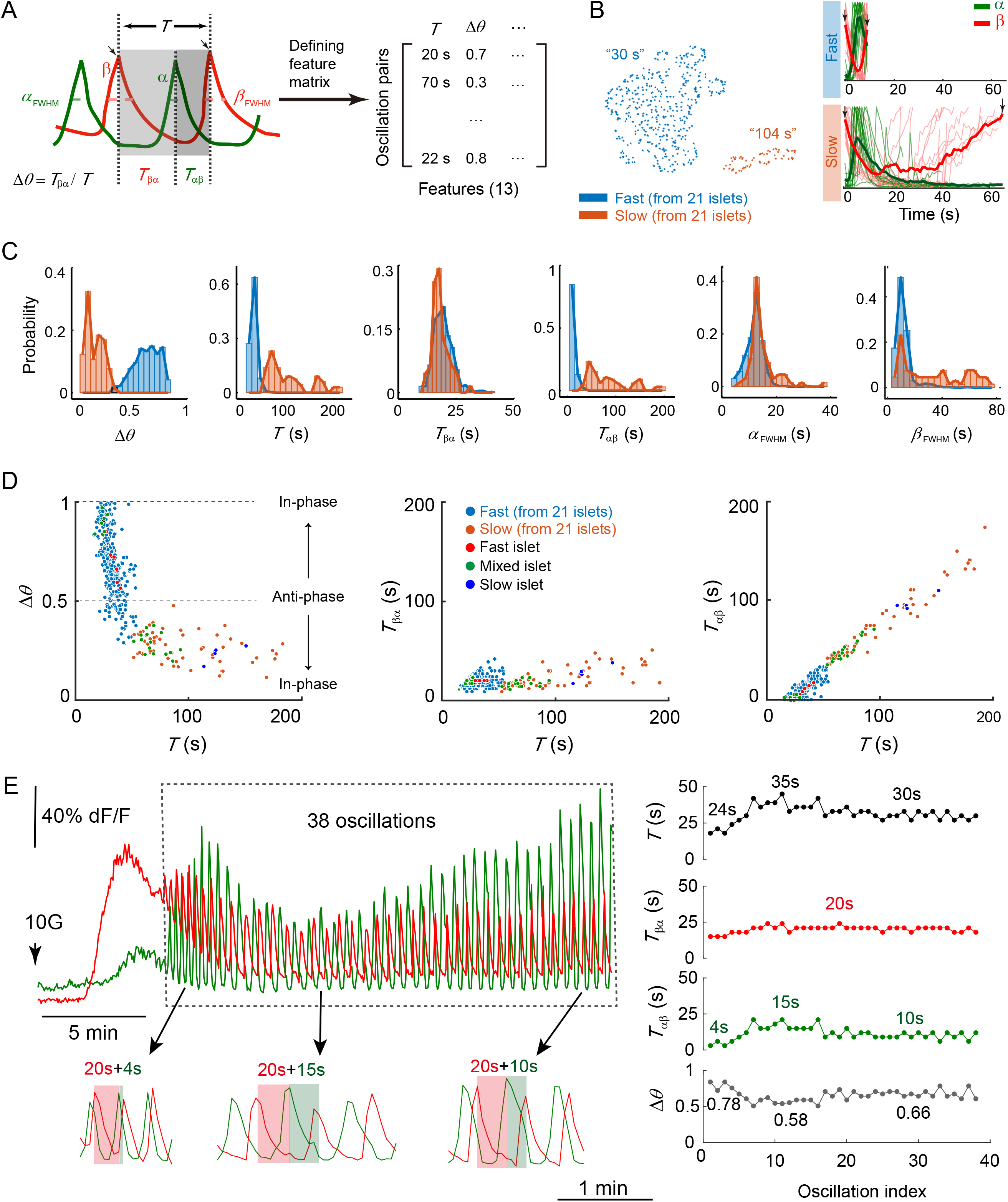
α Cell Activates After β Cell with a Fixed Time Delay While β Cell Activates After α Cell with Variable Time Delay. (A) Left: Each oscillation cycle is defined as the interval between two β cell activations (see Fig. S3 for alternative ways to define the period *T*). Right: 13 features of α-β oscillation cycle are used to construct the feature matrix (columns represent features, rows represent oscillation cycles). (B) Left: Oscillation cycles are separated into two clusters by using UMAP, with the mean oscillation period (*T*) of each cluster shown (n=658 cycles from 21 islets, the fast and slow clusters have 574 and 84 oscillation cycles, respectively). Right: Mean Ca^2+^ traces (bold line) and 15 representative traces (thin line) of fast cluster (top panel) and slow cluster (bottom panel), aligned at the peaks of β cell Ca^2+^ activity (arrows). (C) Histogram of phase difference (Δ*θ*), oscillation period (*T*), waiting time of α and β cells (*T*_βα_ and *T*_αβ_), and Full Width at Half Maximum of α and β cells (*α_FWHM_*, *β_FWHM_*) in fast and slow clusters defined in Fig. S3. (D) Top: Scatter plot of Δ*θ* versus *T.* Dashed lines indicate in-phase and anti-phase α and β oscillations. Middle: Scatter plot of *T*_βα_ versus *T*. Bottom: Scatter plot of *T*_αβ_ versus *T*. The blue and orange dots represent the fast and slow clusters from all 21 islets in (B). The red, green and dark blue dots represent the oscillations from fast, mixed and slow islets shown in Fig. 5D (right panels). (See Fig. S4 for scatter plots of other ways to define the period). (E) Left: Mean Ca^2+^ traces of α and β cells when the glucose stimulation was shift to 10G from 3G. Rectangle box showed 38 α and β phase-locked oscillations. Right: *T*, *T*_βα_, *T*_αβ_ and Δ*θ* of the 38 oscillations.

As *T*, Δ*θ*, *T*_αβ_ and *T*_βα_ were inter-dependent parameters, we further evaluated their relationship using scatter plots (Fig. 4D). It was apparent that *T*_βα_ remained constant across different *T*. In contrast, *T*_αβ_ increased linearly with *T*. These findings strongly suggest a key role of *T*_αβ_ in determining the period (hence the mode) of β cell oscillation. While a short waiting time *T*_αβ_ leads to fast oscillations, long waiting time *T*_αβ_ confers slow ones. Given the α cells displayed a fixed delay of ~20 s to the β cells, the phase-shift between α and β cells varied with the delay of β cells. For example, when the β cells had little delay (~0 s) to the α cells (Fig. 4D, *T*_βα_=~20 s, *T*_αβ_=~0 s, *T*=~20 s), the α and β cells were nearly in-phase (Δθ~=1). When the β cell’s delay was around ~20 s (*T*_βα_~=*T*_αβ_=20 s, *T*=~40 s), the α and β cells were nearly anti-phase (Δθ~=0.5). Finally, when the β cells waited much longer than the α cells (*T*_βα_~=20 s, *T*_αβ_~=180 s, *T*=~200 s), the α and β cells appeared nearly in-phase again but with Ca^2+^ transient much differed from the first case (Δθ~=0.1). Note the apparent order of α and β cell activation depended on the relative value of *T*_αβ_ and *T*_βα_. The fast oscillation might show a near in-phase locking with α ahead of β cell due to a shorter *T*_αβ_ than *T*_βα_, and the slow oscillation might also show a near in-phase locking but with β ahead of α cell due to a shorter *T*_βα_ than *T*_αβ_ (Figs. 3B, 3C and 3F). These relationships hold for various oscillation modes as indicated by the different colors in Fig. 4D.

Interestingly, sometimes we observed that the phase shift between α and β cells underwent an initial transient period during which it changes in time before stabilizing at a steady value. This was usually due to a changing delay of β cell activation after α cell activation. As it is shown in Fig. 4E, while *T*_βα_ remained stable at 20 s during the whole process, *T*_αβ_ started from 4 s, extended to 15 s in the next ten oscillations, and finally stabilized at 10 s. Correspondingly, α and β cells started from a phase shift Δθ=0.78 (*T*_βα_=20 s, *T*_αβ_=4 s) at first, then gradually established a phase shift Δθ=0.58 (*T*_βα_=20 s, *T*_αβ_ =15 s), before finally stabilized at Δθ=0.66 (*T*_βα_=20 s, *T*_αβ_=10 s) (Table S1, n=3 islets). This data corroborated our analysis conducted in multiple islets and pointed to the possibility of dynamic changes in the interactions between α and β cells in the same islet. Because isolated single β cells displayed only slow oscillation with a period of about 6 minutes (Figs. S5A and S5B), we speculate that increased stimulatory effect from α cells to β cells may push β cells to oscillate in the fast mode.

### Mathematical Modeling

We observed that islets show heterogeneous yet intrinsic oscillation patterns under high glucose stimulation. To better understand the origin of various oscillation modes and the factors controlling them, we developed a mathematical model incorporating interactions between α and β cells.

#### α-β Phase Oscillator Model

Given the fact that both α cells and the β cells in an islet were globally synchronized respectively, we simplified the islet as a model of two coupled “cells” - an α cell and a β cell (Fig. 5A). The oscillation of each cell was described by a phase variable *θ*, which was driven by an intrinsic force and a paracrine force (Fig. 5B, see Methods). The intrinsic term corresponded to the oscillation frequency of the single isolated cells. As shown by previous and our studies, single β cells oscillate with a period ~3-6 minutes, and single α cells oscillate with a period ~30-60 seconds (Fig. S5). The paracrine term consisted of three parts: *f*_s_(*θ*) represented the hormone secretion, *f*_rα_(θ) represented the paracrine inhibition of α cells by β cells and *f*_rβ_(*θ*) represented the paracrine stimulation of β cells by α cells. The main results of the model were insensitive to the choice of the specific forms of *f*_s_(*θ*), *f*_rα_(*θ*) and *f*_rβ_(*θ*) as long as they were periodic functions resembling the general characteristic of the biology (see Figs. S6 and S7 for details). The coefficients *K*_αβ_ and *K*_βα_ represented the coupling strengths between the α and β cells. Note that this is a two-phase model without providing any information on the amplitude of the oscillation. Its behavior can be characterized by the “winding number” defined as the asymptotic ratio of the two phases *w=θ*_α_*/θ*_β_ (Fig. 5C).

**Fig. 5.**
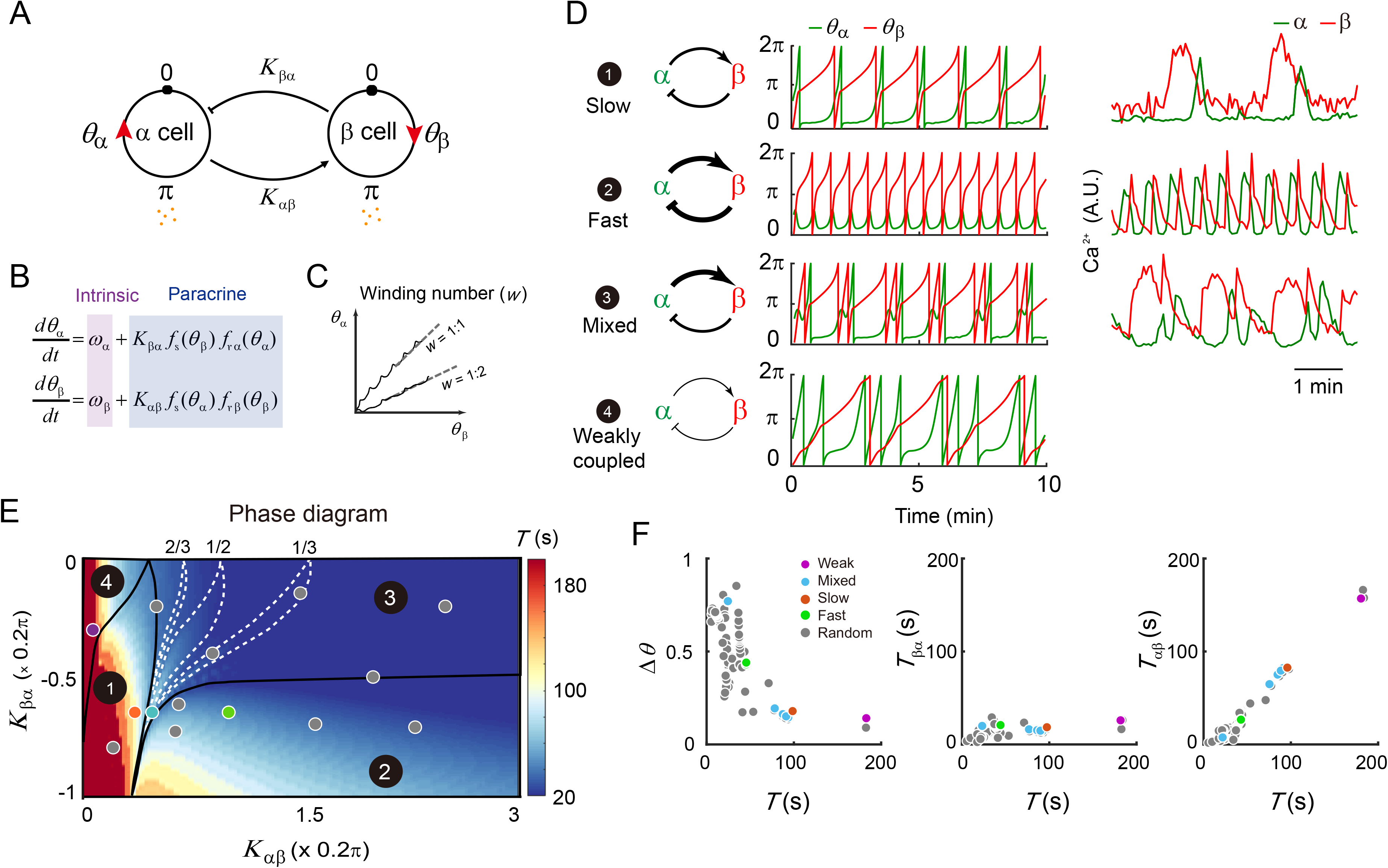
Mathematical Model of Islet α and β Cells. (A) Two-cell islet model. The state of each cell is described by its phase θ. Cell secretes hormone at phase *π*. The hormone secreted by α cell stimulates β cell (with strength *K*_αβ_), and the effect of β cell activation inhibits α cell (with strength *K*_βα_). (B) Equations for phase dynamics of α and β cells. The phase of each cell is influenced by its own intrinsic frequency and the paracrine stimulation/inhibition. Function *f_s_*(*θ*) describes the hormone secretion and function *f_r_*(*θ*) describes the cell’s response to the hormone. (C) Illustration of winding number (*w*) definition. Two examples are shown. Solid lines are the actual trajectories of two solutions of the equations in (B), and dotted lines are straight lines representing the asymptotic limits of the trajectories that define the winding number. (D) Left panel: Schematic of α and β cells’ interaction strengths. Four cases from the four regions in are shown. Middle panel: Example traces of *θ*_α_ and *θ*_β_ in the four cases, respectively. The parameters used in these examples are indicated by the four colored dots in (E). Right panel: Corresponding examples of Ca^2+^ traces found experimentally for the first three cases. (E) The phase diagram of the system. Depending on the two coupling constants *K*_αβ_ and *K*_βα_, the oscillatory behavior of the two cells falls into one of the four phase-locked regions, characterized by the winding number *w*. In region 3, any rational winding number *w*<1 has a stable phase-locking region. For clarity, only the phase-locking regions for lower-order rational numbers (1/3, 1/2 and 2/3) are shown. Color bar codes the period of the oscillation. Except for the four colored dots, 7 randomly selected grey dots are also shown. (F) Scatter plots of Δ*θ*, *T*_βα_ and *T*_αβ_ *versus T*. Each dot represents one oscillation cycle. The color of the dot indicates the parameters used in the simulation, which are shown in the phase diagram (E) with the same color.

#### Slow, Fast, and Mixed Oscillations

By adjusting the coupling strengths (*K*_αβ_ and *K*_βα_) between the α and β cells, our model displayed all three types of the oscillation behaviors observed in experiments (cases 1-3, Fig. 5D). When α cell weakly stimulated β cell (small *K*_αβ_), the model islet generally showed slow oscillations. When α cell and β cell were strongly coupled with each other (large *K*_βα_ and *K*_αβ_), the model islet generally showed fast oscillations. When α cell strongly stimulated β cell and β cell weakly inhibited α cell (large *K*_αβ_ and small *K*_βα_), the model islet generally showed mixed oscillations. With very weak coupling between α and β cells, the model displayed an oscillation behavior similar to uncoupled single cells but not islet experiments (Fig. 5D, case 4). Further quantification found the phase difference Δ*θ* and the period *T* displayed an inverse proportional relationship (Fig. 5F, left panel), which was because of a constant waiting time of the α cell regardless of the oscillation modes (Fig. 5F, middle panel), similar to the experimental data (Fig. 4D). The coupling coefficients changed only the waiting time of the β cell, and further determined the period (Fig. 5F, right panel).

#### Phase Locking Between α and β Cells

We next analyzed the model’s behavior by systematically varying the coupling strengths between the α and β cells. We found that the α and β cells were generally phase-locked, i.e., their phases were dependent on each other with a fixed relationship characterized by the winding number *w* (Fig. 5C). By plotting the winding number’s dependence on the coupling strengths, we obtained the phase space of the system of two coupled oscillators (Fig. 5E). It formed a two-dimensional Arnold tongues^48^, which could be separated into four regions (Fig. 5E, see Methods). In region 1, the α cell and the β cell are locked on the *w*=1/1 mode. That is, when *θ*_β_ finishes one cycle ([0,2π]), *θ*_α_ will also finish one cycle. An example of this oscillation mode is shown in Fig. 5D (upper panel). In region 2, the two cells are locked in the mode *w*=0/1. That is, when *θ*_β_ finishes one cycle ([0,2π]), *θ*_α_ cannot finish a full cycle before being pushed back (Fig. 5D (middle-upper panel)). Here the strong stimulation from α to β induces a fast oscillation frequency, while the strong repression from β to α prevents the α cell from finishing a full cycle every time it is activated. In region 3, the two cells are locked with 0<*w*<1. In particular, there exist *w*=*m*/*n* modes, where *m*<*n* are both integers. In a mode with *w*=*m*/*n*, when *θ*_β_ finishes *n* cycles ([0,2*n*π]), *θ*_α_ will finish *m* cycle(s) ([0, 2*m*π]). In the example with *w*=1/2 shown in Fig. 5D (middle-lower panel), while each activation of the β cell can finish one full cycle, the first activation of α cell cannot finish a full cycle before being pushed back, and only the second activation can finish a full cycle. Although only a few modes are shown in region 3 for clarity, it can be proved rigorously that region 3 contains all the modes, with *w* being a rational number between 0 and 1 (unpublished). In region 4, the two phases are locked with *w*>1, which means that α cell will finish more cycles than β cell. An example of *w*=3/1 is shown in Fig. 5D (lower panel). In this region, α and β cells couple weakly. At the upper left corner *K*_αβ_=*K*_βα_=0, α and β cells completely decouple and beat on their intrinsic frequencies. Note that while *w* jumps discontinuously with continuously varying parameters, the average period of β cell oscillation changes smoothly (Fig. 5E, heat map). Thus, the paracrine interaction between α and β cells offers robust and tunable oscillation patterns and periods.

#### Model Prediction and Verification

A prediction of the model was that the oscillation period may be tuned with the strengths of paracrine interaction, depending on the original position of the islet system in the phase space (Figs. 5E, heat map, 6B, S8E and S9F). In particular, increasing the activation from α cell to β cell (*K*_αβ_) could increase the oscillation period, especially in islets of slow oscillations (Fig. 5E, heat map). We applied glucagon (100 nM) to the islets showing fast and slow oscillations (Figs. 6A and S8A). While adding glucagon did not affect fast oscillating islets, it switched islets harboring slow oscillations into fast ones. According to our model, the change of oscillation period was due to stronger paracrine interactions that reduced the waiting time *T*_αβ_ without affecting the waiting time *T*_βα_, which faithfully recapitulated the experimental data (Figs. 6B-D and S8B-E). On the other hand, the model predicted that reducing the effect of glucagon may lead to more autonomous cellular regulation and slow oscillations (Figs. 6B, 6F and S9F). Indeed, by combining insulin and the GCGR and GLP-1R antagonists (MK0893 (MK) and Exendin 9-39 (Ex9)) to inhibit glucagon secretion and its downstream target^30,31^, ~50% of the fast oscillatory islets switched to slow oscillations (Fig. 6A). The change of modes reversed back when the inhibitory agents were removed (Fig. S9A). Islets’ fast-to-slow mode switching critically relied on the activation level of glucagon’s downstream target. A weaker glucagon receptor antagonist combination prolonged the islet oscillation period without inducing fast-to-slow mode switching (Figs. S9B and S9C). The β-cell-specific GCGR knockout mice (*Ins1-cre;Gcgr^f/f^*)^49^ had fewer fast oscillation islets (Fig. S9D). And ~50% of the fast oscillation *Ins1-cre;Gcgr^f/f^* islets turned into slow oscillations with the weaker glucagon receptor antagonist combination (Fig. S9E).

**Fig. 6.**
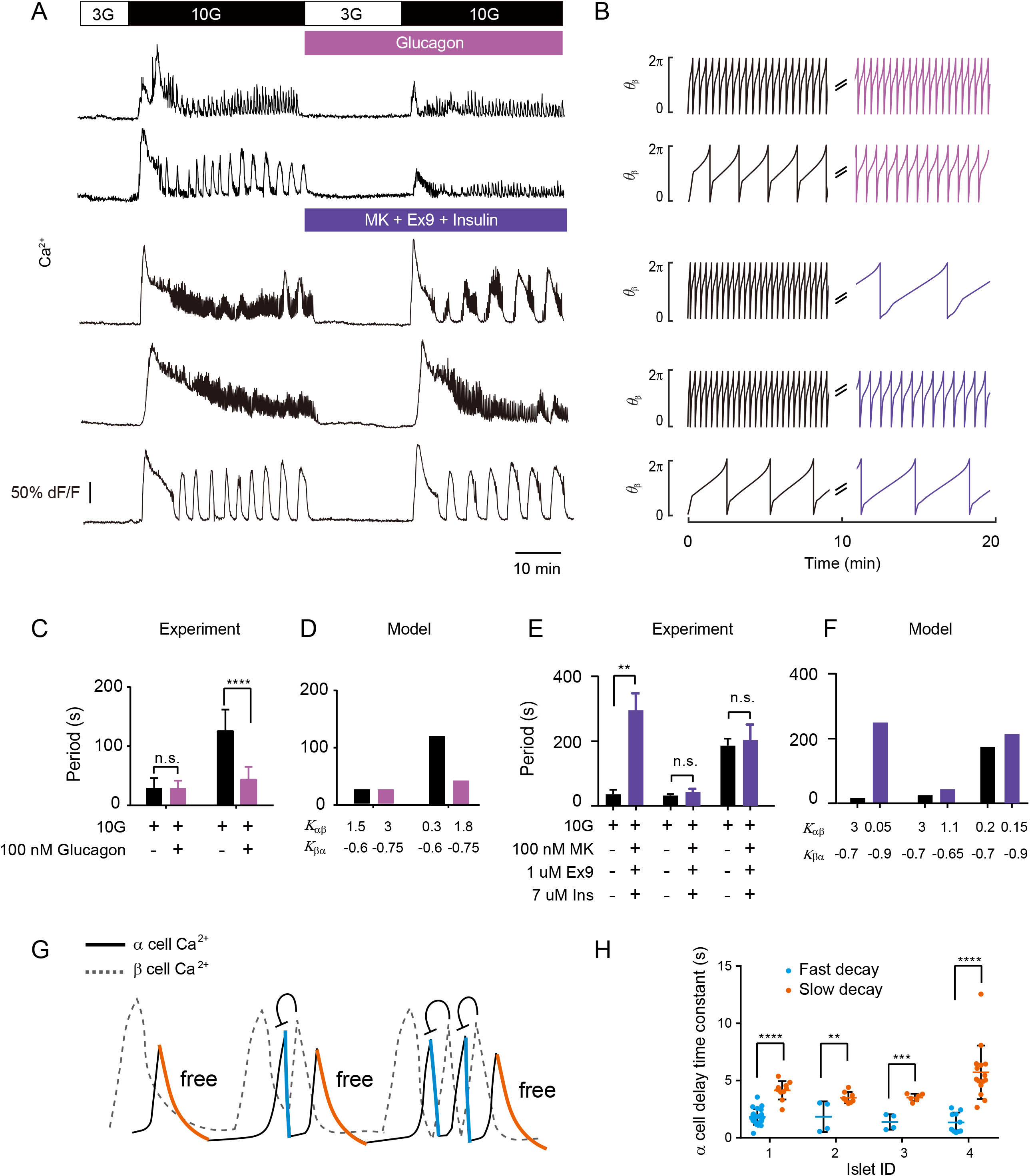
Model Predictions Verified by Experiments. (A) Top two rows: Representative recordings of β cell Ca^2+^ signal for fast and slow islets in *Glu-Cre^+^;GCaMP6f^f/+^*; *Ins2*-*RCaMP1.07* mice. (See Fig. S8A for more examples and with both α and β signals). The stimulation used in the experiment is shown above: 3G (10 min), 10G (40 min), 3G + 100nM glucagon (20 min) and 10G + 100nM glucagon (35 min). Bottom three rows: Representative recordings of β cell Ca^2+^ signal for fast and slow islets in *Ins^+/-^;GCaMP6f^f/+^* mice. The stimulation is 3G (10 min), 10G (40 min), 3G + 100nM MK (MK0893) + 1uM Ex9 (Exendin (9-39)) + 7uM insulin (20 min) and 10G + 100nM MK + 1uM Ex9 + 7uM insulin (35 min). (B) Modeling results for increasing and decreasing *K*_αβ_ and *Kβα* on fast and slow islets. First row: *K*_αβ_=1.5, *Kβα*=−0.6 for 0-10 min and *K*_αβ_=3, *Kβα*=−0.75 for 10-20 min. Second row: *K*_αβ_=0.3, *Kβα*=−0.6 for 0-10 min and *K*_αβ_=1.8, *Kβα*=−0.75 for 10-20 min. Third row: *K*_αβ_=3, *Kβα*=−0.7 for 0-10 min and *K*_αβ_=0.05, *Kβα*=−0.9 for 10-20 min. Forth row: *K*_αβ_=3, *Kβα*=−0.7 for 0-10 min and *K*_αβ_=1.1, *Kβα*=−0.65 for 10-20 min. Fifth row: *K*_αβ_=0.2, *Kβα*=−0.7 for 0-10 min and *K*_αβ_=0.15, *Kβα*=−0.9 for 10-20 min. (C) Mean Ca^2+^ oscillation period from experiments under 10G stimulation without and with glucagon treatment (n=4 fast-to-fast and 3 slow-to-fast islets). (D) Mean period from the model in the top two rows of (B). (E) Mean Ca^2+^ oscillation period from experiment under 10G stimulation without and with the combination of insulin, MK and Ex9 (n=4 fast-to-slow, 5 fast-to-fast and 5 slow-to-slow islets). (F) Mean period from the model in the bottom three rows of (B). (G) Schematic of Ca^2+^ trace in islets of mix oscillations. The solid line shows α cell Ca^2+^ activity and the dashed line β cell. The α cell transients immediately preceding a β cell activation decay faster (colored orange) due to the β cell’s repression. The α cell transients away from β cell activation would decay with their own intrinsic dynamics and thus slower (colored blue). (H) The decay time constants (see Fig. S3 for the definition) for the two kinds of α cell transients described in (E) in mixed oscillations. They fall into two groups: fast (yellow) and slow (blue). Data from 4 islets are shown. Bars represent mean ± SEM (standard error of mean). n.s. p>0.1, *p<0.05, **p<0.01, ***p<0.001, ****p<0.0001. Statistical comparisons are conducted using unpaired t test.

Finally, our model also predicted that in mixed oscillation modes, decay times of Ca^2+^ transients in α cells were different - only the last Ca^2+^ transient in each cluster of cycles of mixed oscillation was independent of β cells, while all other ones were repressed by β cells and should descend faster (Fig. 6G). By analyzing the Ca^2+^ traces of islets with mixed oscillation modes, we confirmed that the decay times of α cell transients fell into two groups: the Ca^2+^ transients in α cells just preceding an uprising of β cell activation decayed faster. In contrast, the ones posterior to β cell transients exhibited a significantly slower decay (Fig. 6H). Overall, the agreement between the model and experiment highlights the importance of α-β interactions in shaping up the oscillation modes.

## DISCUSSION

In this study, we developed a microfluidic device that enabled stable and repeatable long-term imaging of Ca^2+^ activities of islets at single-cell resolution. Despite the apparent heterogeneity in Ca^2+^ activities across different islets, individual islets exhibited their own spatial and temporal patterns of Ca^2+^ oscillations that were repeatable under multiple rounds of glucose stimulation. This suggests that the oscillation mode results from some intrinsic properties of the islet, possibly correlated with different cell types and their spatial distributions.

By using the *Glu-Cre^+^; GCaMP6f^f/+^; Ins2-RCaMP1.07* transgenic mice, we discovered that the α and β cells were globally phase-locked to various oscillation modes. The PiggyBac approach led to the sparse labeling of β cells – it enabled a clear separation of α and β cell Ca^2+^ dynamics. Besides α and β cells, pancreatic islet δ cells have recently received increasing attention in glucose regulation^50^, which are not included in the current model. δ cells are connected to β cells by gap junctions and exhibit Ca^2+^ activities similar to β cells^51^. It releases somatostatin to strongly inhibit both α and β cells. Further study about the δ cell Ca^2+^ dynamics and simultaneous α, β, and δ cell Ca^2+^ imaging would be important to understand the role of δ cell in tuning α and β phase-locking patterns. In the current mathematical model, the role of δ cells in regulating the oscillation modes was lump-summed together with that of β cells. It is essential to differentiate the two types of cells in future modeling.

A key finding in our study is that the time delay for α cells’ activation following the activation of β cells (*T*_βα_) was invariant, regardless of the islet-to-islet variations in oscillation frequencies and modes. This observation of invariant *T*_βα_ echoed nicely with the ~20 seconds recovery time of α cells from the relief of optogenetic activation of β and δ cells^51^. Therefore, *T*_βα_ is likely to be determined by one or more secretin released by β and δ cells, including insulin, Zn^2+^, ATP, GABA, and somatostatin^52^.

In contrast, what varied in different islets under different conditions was the time delay of β cells’ activation following that of α cells (*T*_αβ_). Our and previous studies suggest that glucagon is a contributing factor in tuning the oscillation mode^13^. Our work showed that increased glucagon level tunes on the oscillation modes by reducing the waiting time of β cells (*T*_αβ_). Glucagon may elevate cytosolic Ca^2+^ concentration and increase oscillation frequency, both through cAMP-dependent^53,54^ and independent pathways^55–58^. On the other hand, blocking glucagon receptors with MK0893 and Exendin 9-39 could not fully slow down WT islets. We speculate that this may be because pharmacological inhibitors cannot completely block the endogenous glucagon function of pancreatic islets. Consistent with this, MK0893 and Exendin 9-39 induced fast-to-slow mode switching in the *Ins1-cre;Gcgr^f/f^* islets, and a combination of insulin, MK0893 and Exendin 9-39 was able to turn fast oscillations into slow ones. The specific molecular mechanism for this synergy effect needs future investigation.

The fixed α cell delay and glucagon-tuned β cell delay imply that the value of the phase shift depends on the oscillation frequency (Fig. S10). Islet β cell Ca^2+^ oscillation modes are influenced by two regulatory processes – the fast-paracrine stimulation and the slow intrinsic activation. Once α cells release a large amount of glucagon, their stimulatory effects on β cells would significantly reduce the waiting time for β cell activation and the islet would oscillate with a frequency much faster than β cells’ intrinsic frequencies (α paracrine dominant). If α cells fail to release enough glucagon, or the downstream effects of glucagon are inhibited, β cells in the islet would oscillate with a frequency close to their intrinsic ones (β intrinsic dominant). Indeed, the α and β cells appeared nearly in-phase for both fast oscillations with periods close to 20s and slow oscillations with period ~180s. There is a continuum between the two extreme cases. E.g., for oscillations with period ~40s, α and β cells appeared nearly anti-phase.

In our study, both experimental observations and model simulations showed a robust phase-locking phenomenon between α and β cells. It is known that faster islet Ca^2+^ oscillations display more regular oscillation patterns^13,59^. In light of our findings, the increased regularity may come from the increased stability in phase-locking: more rapid oscillation implies more robust activation from α to β cells and thus a tighter regulation. The Ca^2+^ oscillation of the two types of cells can phase-lock to a variety of modes determined by the paracrine interactions, which not only have different oscillation frequencies, but also display a range of other quantitative features such as the winding number and the ratio of the half-widths for α and β transients. Phase-locking to different frequencies and modes could ensure a stable and tunable secretion of insulin and glucagon. A variety of β cell Ca^2+^ oscillation modes were observed *in vivo*, including those of fast ones^60–62^. Further studies combining islet Ca^2+^ imaging with real-time detection of α and β cell secretion are needed to investigate the physiological roles of the phase-locking and its dependency on paracrine interactions.

Previous studies tried to explain the distinct Ca^2+^ oscillation modes in intact islets with single beta-cell models, which did not consider the contribution from other cell types in islets^15–18^. Our model emphasized how paracrine interactions may play important roles in various islet Ca^2+^ oscillation modes. It is conceivable that both intrinsic properties of single cells and interactions among different cell types may contribute to the regulation of islet Ca^2+^ oscillation modes. Further investigation using pseudo-islets with varying compositions of α and β cells may help to differentiate the roles of intrinsic and paracrine contributions^63^.

## ACKNOWLEDGMENTS

We thank Erik Gylfe, Anders Tengholm, Yanmei Liu, Louis Tao and Alexander Valentin Nielsen for helpful discussions. We thank Chunxiong Luo and Shujing Wang for their help with the microfluidic chip design. We thank Daniel Tang and Iain Bruce for manuscript editing. We thank Xiaowei Chen for sharing mice lines. This work was supported by grants from the Chinese Ministry of Science and Technology (2015CB910300), National Natural Science Foundation of China (81925022, 92054301), The National Key Research and Development Program of China (SQ2016YFJC040028, 2018YFA0900700), NSFC Innovation group projects (31821091) and the Beijing Natural Science Foundation (L172003, 7152079, 5194026).

## AUTHOR CONTRIBUTIONS

C.T. and L.C. conceived and supervised the study, and wrote the manuscript.

H.R. led the project, designed experiments, carried out data analysis and mathematical modeling, and wrote the manuscript.

Y.L. designed and manufactured the microfluidic chip, designed and performed experiments, and wrote the manuscript.

C.H. designed and performed experiments, and wrote the manuscript.

X.Y. contributed to Figure preparation and supervised experiments, Y.Y and K.S. contributed to the mathematical modeling, B.S. and S.W. contributed to data analysis.

